# Simulated Language Acquisition in a Biologically Realistic Model of the Brain

**DOI:** 10.1101/2025.07.15.664996

**Authors:** Daniel Mitropolsky, Christos H. Papadimitriou

## Abstract

Despite tremendous progress in neuroscience, we do not have a compelling narrative for the precise way whereby the spiking of neurons in our brain results in high-level cognitive phenomena such as planning and language. Nemo is a simple mathematical formulation of six basic and broadly accepted principles of neuroscience: excitatory neurons, brain areas, random synapses, Hebbian plasticity, local inhibition, and inter-area inhibition. We implement with Nemo a simulated neuromorphic system, which is capable of basic language acquisition: starting from a tabula rasa, the system learns, in any language, the semantics of words, their syntactic role (verb versus noun), the word order of the language, and the ability to generate novel sentences, all through the exposure to a number of grounded sentences in the same language that is similar to language acquisition by humans. We discuss several possible extensions and implications of this result.

## 1 Introduction

It is beyond doubt that cognitive phenomena such as language, reasoning, and planning are the direct product of the activity of neurons and synapses, and yet there is no existing overarching theory that explains exactly how this could be done. In the words of Richard Axel (1), *“we do not have a logic for the transformation of neural activity into thought and action. I view discerning [this] logic as the most important future direction of neuroscience*.*”*

We introduce such a minimalistic neural model of the brain, which we call Nemo (see (27) for previous work on Nemo, and Figure 1 for a brief exposition), a simple mathematical formulation of six basic and uncontroversial ideas in neuroscience: excitatory neurons, brain areas, random synaptic connectivity, local inhibition in each area, Hebbian plasticity, and interarea inhibition. Importantly, Nemo can be simulated very efficiently at the scale of tens of millions of neurons and trillions of synapses. To test whether Nemo qualifies as the abstraction sought in Axel’s statement, we design a Nemo system that can carry out what is arguably the greatest achievement of the animal brain, namely *natural language acquisition*. In particular, we create a tabula rasa Nemo system that learns, for any natural language (a) semantic representations of concrete nouns and verbs; (b) basic syntactic characteristics such as the part of speech for each word; (c) the language’s word order, in several moods, and (d) to generate novel sentences — all through exposure to a modest amount of grounded language.

**Figure 1.**
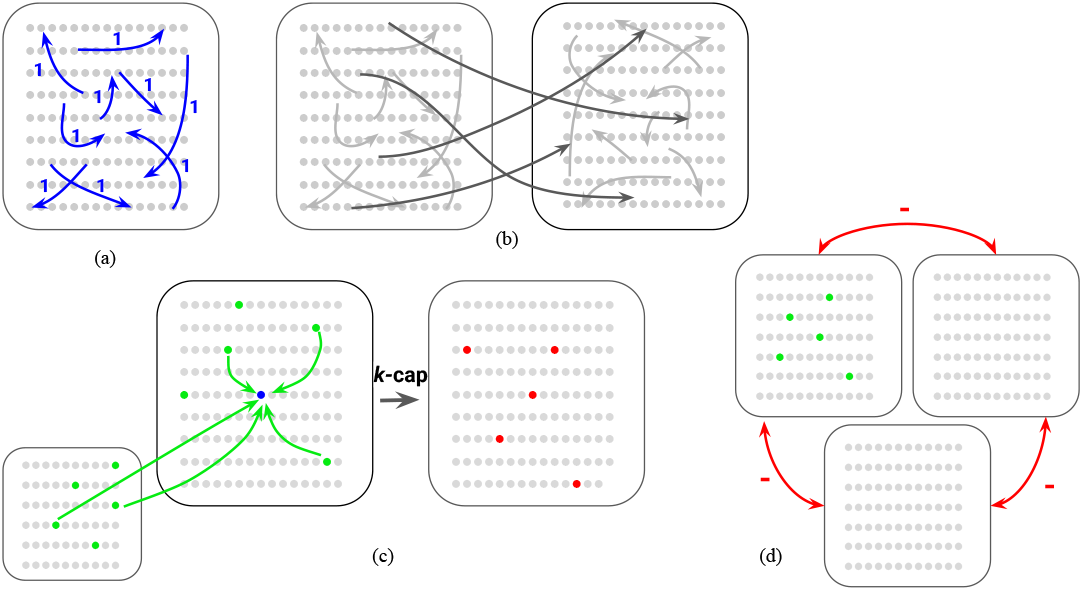
The Nemo model (see (27) for a detailed exposition): (a) the brain is a set of brain *areas*, each containing *n* excitatory neurons (typically 10^6^). Every synapse between neurons is sampled with probability *p* (typically 0.01) and initialized with weight 1. (b) an ordered pair of areas may be connected by a *fiber*, i.e. with randomly sampled synapses. Normally only a few of all possible fibers between areas are present, defining the top level of the system’s architecture. (c) Nemo is a dynamical system defined by *firing*. At each discrete time step, only *k* neurons in each area fire — a selection that models local inhibition in the area. The parameter *k* is much smaller than *n* (typically about 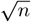). Of the *n* neurons in an area, the *k* that fire are those that received the *largest weighted synaptic input* from the previous time-step; *this primitive of the* Nemo *model is called k*-*cap*. There is also *Hebbian plasticity:* for a synapse from neuron *j* to *i* firing in succession in this order, the synaptic weight *wj,i* is multiplied by (1 + *β*); the positive plasticity parameter *β* is typically 0.05. (d) Additionally, two or more areas can be in *mutual inhibition*, in which case there is firing only in the area that receives the greatest total synaptic input from the previous step. Notice that, so far, synaptic weights could increase with no limit, and this can be alleviated by weight normalization.

A complete account of language acquisition would also include the acquisition of speech sounds and word phonetics, parts-of-speech other than nouns and verbs, including functional categories of words such as “the,” abstract words, and syntax beyond basic word-order. Regarding speech sounds, we adopt a convenient input/output convention: words are input to the device and output from it as tokens, through the activation of phonetic word representations assumed to be already in place. That is, we bypass the phase of establishing linguistic input/output in order to focus on the aspects of language that are exclusive characteristics of humans — it is known that primates can understand and generate individual signed words (29). In the next section, we present our basic language acquisition system implementing the core statistical learning tasks (a)–(d) above in two phases: first the learning of word semantics and part of speech (a–b) and then the learning of word order and generation (c–d). Then we discuss several extensions and variants of the basic system that address some of the shortcomings mentioned above, and show that our results are in good agreement with what is known about language acquisition by children.

In our experiments, the learning device is exposed to *whole grounded sentences* in any fixed natural language (we use several languages in our experiments). By “grounded” we mean that each sentence is presented, while representations corresponding to the sentence are active in the sensory and motor cortex. That is, when the sentence “the dog runs” is input, we assume that in the premotor cortex the mirror cells (19, 9) for “run,” as well as the generic “dog” representation in the inferotemporal visual cortex (18), are firing. In a very interesting precursor to this work (30), see also (13), it was shown that concrete nouns, verbs and their meanings can be learned in a different biologically realistic model of the brain. However, that work was about the easier problem of learning *isolated words*, without syntactic context — it is known that nonhuman primates can achieve this level of language learning, see, for example (29).

Note that, besides addressing the quest for Axel’s “logic,” our work is about another fundamental question: *What is the biological basis of language?* Is language the fruit of unique human characteristics at the genetic, molecular, and/or neural level, such as the FOXP2 gene (once called “the language gene”) or the complex dendritic potentials of human neurons (15, 6)? Our work is evidence that a language system can be built on top of the mammalian brain, through well accepted neuroscience — albeit through the evolution of a system of brain areas and fibers whose level of complexity may not exist elsewhere in the animal brain. The specific neural architecture and algorithms we propose here can be seen as a comprehensive, neurobiologically plausible *hypothesis* for the human language system.

## 2 Procedures and Results

### 2.1 Learning word semantics

See Figure 2 for the Nemo architecture (areas and fibers) for learning word semantics. As explained before, we bypass the stage of acquiring speech sounds through an *input/output convention:* words are input to the device through the activation of word representations in a brain area Phon (Figure 2), anticipated in the literature (16, 26), containing phonetic knowledge for recognizing and articulating words. The *lexicon* consists of two areas, Lex_1_ and Lex_2_, where assembly representation for each word in the lexicon is formed: in Lex_1_ for nouns and in Lex_2_ for verbs – this separation is also anticipated in much of the literature (25, 31, 20), even though others differ.

**Figure 2.**
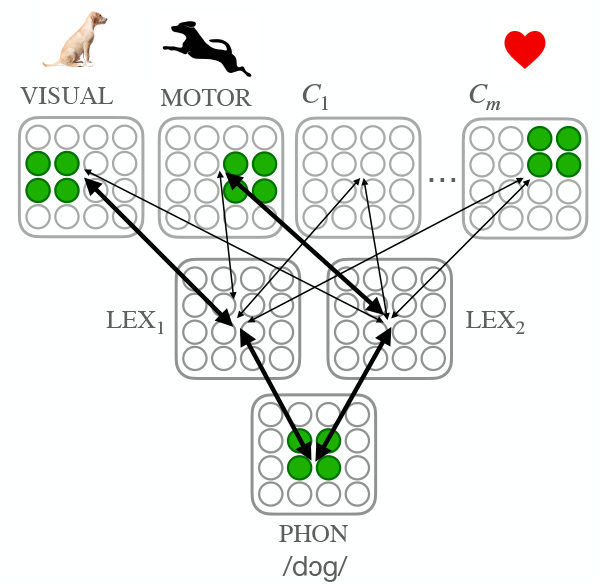
The Nemo system for learning concrete nouns and verbs. Phon is an input area which is initialized with one representation, for each word, of the phonological code of the word. The semantic areas Visual, Motor and *C*_1_, …, *C*_*m*_ are also input areas, with visual representations, motor representations, and other perceptual representations (e.g. olfactory, emotional, etc). Lex_1_ and Lex_2_ are the areas where the nouns and verbs, respectively, in the language will be learned. Lex_1_ has high connectivity to Visual and Lex_2_ to Motor, and lower connectivity to the other semantic areas. The figure shows the firing pattern in the input areas when input the sentence “dog jumps,” at the moment “dog” is heard.

The system also has a number of *semantic areas*, sensorimotor and other cortical areas that feed into the language system, whose number varies in our experiment from 2 to 12. — the existence of such areas was noted, e.g., in (28). Two of these areas, Visual and Motor, have special significance. Like Phon, the semantic areas are initialized to contain assembly representations, here to provide semantic content to words in the lexicon, mirror neurons for concrete verbs and visual representations for concrete nouns, but also olfactory, affective, and other kinds of representations.

#### The word learning experiment

We fix a natural language and *ℓ* concrete nouns and *ℓ* intransitive action verbs (*ℓ* is a parameter that we vary in our experiment). By restricting ourselves to action nouns and concrete verbs we assure that for each word in the lexicon there is a corresponding semantic representation in the semantic areas. The restriction to intransitive verbs is considered only for the present experiment of learning word semantics; transitive verbs will be introduced soon. All sentences in this experiment are of length two: “the cat jumps” and “the dog eats,” where, for the purposes of our experiment, we assume that function words such as “the” are simply ignored; taking into account such words makes the learning task easier. Note that the language can have either SV (subject-verb) principal word order (as in English, Chinese and Swahili) or VS (as in Irish, Modern Standard Arabic, and Tagalog), and our model should succeed in either scenario; this is one of the main challenges of this experiment.

For each word *w* we initialize an assembly Phon[*w*] in Phon. If *w* is a noun, we initialize an assembly in Visual corresponding to the visual representation of the object of *w* in the inferotemporal cortex, and if *w* is a verb, there is a corresponding assembly in Motor — presumably the mirror cells for *w* in the premotor cortex. Finally, for a subset of the remaining *m* context areas, selected at random, we create in each area an assembly corresponding to *w*.

We input randomly generated sentences via a uniform choice of noun and verb. To input a sentence, we input each word individually, in the word order of the language, by firing the corresponding assembly in Phon *τ* times, where *τ* is a parameter of the experiment, while the contextual input of the scene (the assemblies associated with the words in the sentence, both with the noun and the verb) fire throughout the duration of the entire sentence. Note that it is a priori impossible to know which of the two words is the concrete noun, and which is the action verb. However, as we shall see, the interaction between Nemo dynamics and cooccurrence statistics results in the system learning to correctly distinguish between noun and verb.

What does it mean for the system to succeed? Informally, we would like Lex_1_ to become a “noun area,” and Lex_2_ a “verb area,” containing *robust neural representations of each noun and verb* in the lexicon, respectively, and we would also like each of these robust word representations to be somehow aware of the meaning of the corresponding word, as well as aware of their affinity with the corresponding phonetic assembly in Phon. More formally, we require that, after the input of enough sentences, the following properties hold:

##### Property 1: Formation of word assemblies

For each noun, if we fire its representation in Phon, or that in Visual, or both, the resulting *k*-cap in Lex_1_ is stable.

##### Property 2: Formation of the PHON-LEX-VISUAL pathway

The creation of assemblies for each word enables a bidirectional pathway between Phon and the semantic areas via Lex_1_ and Lex_2_: for each noun *w*, if we fire Phon[*w*] for 2 time steps, via Lex_1_ we activate Visual[*w*] and its representation in the contextual areas. Conversely, if we fire Visual[*w*] we retrieve Phon[*w*], and similarly for verbs.

##### Property 3: Classification of Nouns vs. Verbs

While robust assemblies for nouns are formed in Lex_1_ and for verbs in Lex_2_, *nothing akin* to a stable representation exists for nouns in Lex_2_, nor for verbs in Lex_1_. Concretely, firing Phon[*w*] for a noun (resp. verb) into Lex_2_ (resp. Lex_1_) activates an unstable set of neurons such that after one time step, less than 50% of the same neurons fire (and firing back from this set into Phon does not activate Phon[*w*]).

### 2.2 Results of the basic word learning experiment

We experiment with different word orders and lexicon sizes, and we input to the system two-word sentences *until all three Properties 1-3 above hold* thus determining the minimum number of sentences needed for this, as a function of the number *ℓ* of nouns and verbs. Recall that the amount of linguistic input required for language to be learned is of central interest in linguistics, for example, in relation to the “poverty of stimulus” argument (5); our experiment offers an opportunity to actually quantify what “modest linguistic input” means in this debate. Indeed, the required number of sentences appears to *grow as a small linear function* of the number of words in the lexicon (Figure 3a).

**Figure 3.**
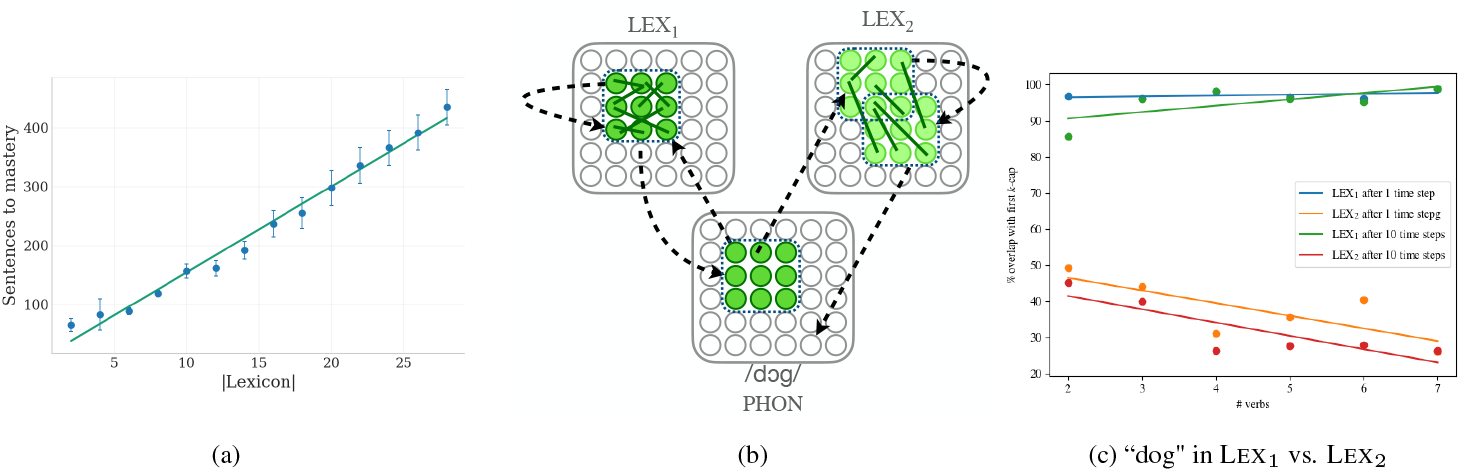
Results of the word learning experiment. In (a) the learning simulation is performed for varying sizes of the lexicon until all of Properties 1-3 hold for the first time for every word, revealing a linear trend (b) illustrates the formation of a stable assembly (dark green) representing “dog” in Lex_1_; in contrast, an unstable and loosely connected set of neurons (pale green) is formed in Lex_2_. (c) the same stability/instability pattern is illustrated numerically, for a varying number of verbs with which “dog” is paired in training sentences. Stability is measured in terms of the overlap of the *k*-cap obtained at the first firing of Phon[*dog*] and that of subsequent firings (one step and ten steps later).

### 2.3 Further experiments and comparisons

We have repeated our word learning experiments in order to explore alternative and/or more realistic variants of our model. We have also compared our model’s performance with data from the language acquisition literature.

First, the neurolinguistic literature is somewhat equivocal on whether verbs and nouns are represented in separate areas in the human brain; accordingly, we repeated the experiment with only one Lex area. It turns out that robust word representations emerge in this case as well, and in fact slightly earlier, with respect to language input, than in the model with two Lex areas; see Figure 6 in SI, as well as Figure 7 which compares acquisition times in the one- and two-LEX versions along with human acquisition times based on (24). For the single-LEX version to succeed, however, the size of the cap in the area Phon (or the synaptic probabilities in the fiber joining Phon with Lex) must be larger — in fact, when these parameters get even larger, the linguistic input required for learning decreases, as shown in Figure 6.

Second, an important part of language acquisition is the exposure of the learner to function words such as pronouns and “the” or, more generally non-content words. It turns out that the model is robust to the presence of such words. Consider the following experiment: In our sentences we present assemblies in PHON, intermingled with the verbs and the nouns, that do *not* have corresponding assemblies in VISUAL, MOTOR, or other contextual areas. When such words are interjected in sentences with reasonably low frequency (e.g. in 75% of sentences), our experiments succeed, and stable representations of such words in the LEX areas fail to form; see Figure 11 in the Supplementary Materials for a description of this experiment. The question of how function words *are* eventually learned and represented, perhaps in other lexical areas, is an interesting open problem. Note that children learn functional categories considerably later than they do concrete nouns and verbs, in terms of the number of exposures (or ELIs, see discussion in next section) (24).

Third, in our experiments so far, the *ℓ* nouns and *ℓ* verbs were sampled uniformly to form input sentences. However, our system works even if the distribution is highly nonuniform, and, in fact, when words are occasionally input individually instead in the context of a sentence, as well as if many more verbs are presented, or many more nouns. In this experiment we track the total number of grounded exposures to a word until a stable assembly for this word is formed. In the case of the 2 LEX areas we also track the number of co-occurrences of a noun with *different* verbs — as we shall see, these are the metrics used in language acquisition research. These exposures can occur over various time scales (e.g. uniformly as in the main experiments of Figures 3 and 6, or all in rapid succession, or spread over time in other ways — see Figures 10 and 9 in the Supplementary Materials for results of learning under such distributions.

#### Comparison with language acquisition data

Between 8 and 20 months toddlers go from producing 0 to 30 words on average — a finding consistent across English, Spanish and Portuguese, see (12). The rate of learning in this period is roughly linear (see Figure 7 overlaying the learning curve of the NEMO model with the average child learning rate based on the model of (24)). A beautiful line of work culminating in the models of (24, 17) has found that, particularly within the first 30 months of life, the moment a toddler begins to produce (or understand) a word is best modeled as a function of the number of *effective learning instances (ELIs)* — contextualized exposures to the given word — with an average of 10-15 ELIs necessary and sufficient to successfully learn a word. This rate is remarkably consistent across languages and part-of-speech (see Figure 5 of (24)). In NEMO, the rate of word-learning depends somewhat on the choices of meta-parameters such as *k* and *β*; however, in both our experiments — with two LEX areas and one LEX area — we find in that our model needs *10-12 contextualized exposures per word to learn a word successfully*.

**Figure 4.**
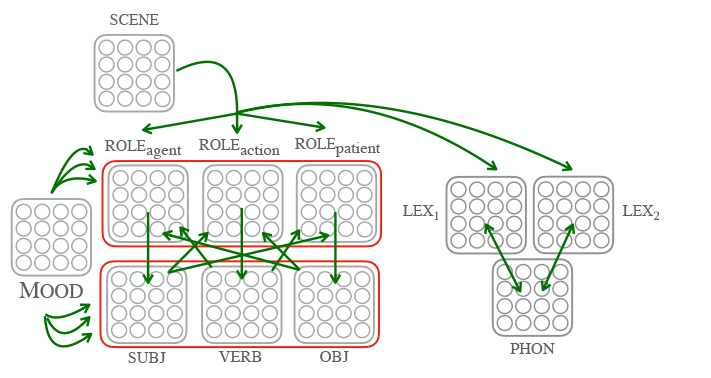
The Nemo system for constituent order learning, as described in Section **??.** Red boxes represent two triples of areas under mutual inhibition.

**Figure 5.**
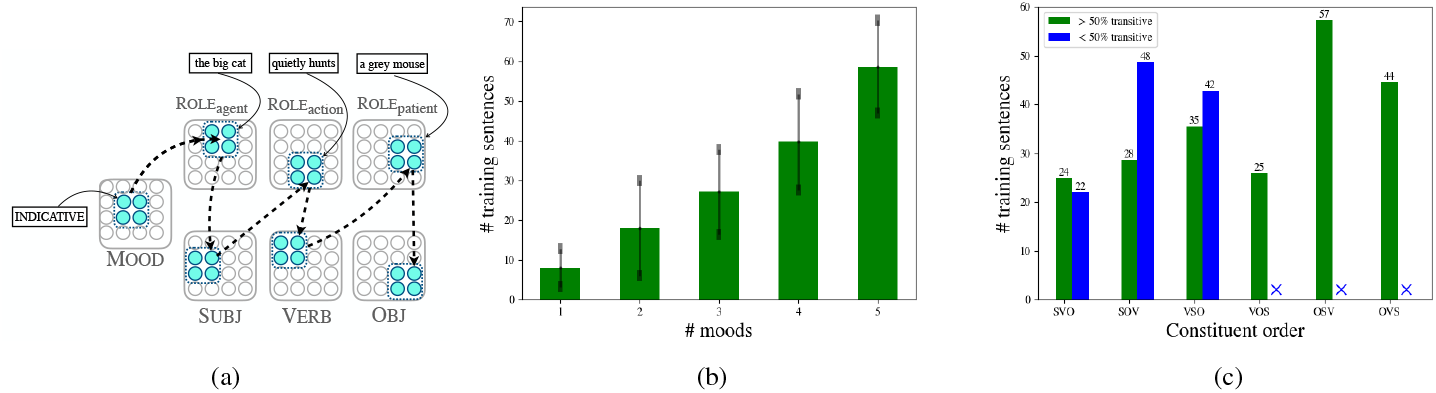
(a) An example of the chain of assemblies that fires in a SVO language after successful constituent-order acquisition. we imagine an expanded system where the assemblies in each Role area can represent whole *phrases*, to suggest how our algorithm for constituent order learning and production is not confined to 3-word sentences. (b) Our system can learn languages with multiple word-orders corresponding to distinct grammatical moods (each mood corresponding to a distinct assembly in the Mood area). (c) We compare the number of sentences needed to learn different constituent orders. We find that significantly more sentences are needed to learn OSV and OVS. If the proportion of transitive sentences is less than half, it is easy to show via rigorous proof that the orders VOS, OSV, or OVS *cannot* be learned by this system even with unbounded input, and so either an altogether different mechanism, or predominance of transitive input sentences must account for the acquisition of syntax in such languages — note that these are in fact the three rarest constituent orders among world languages.

**Figure 6.**
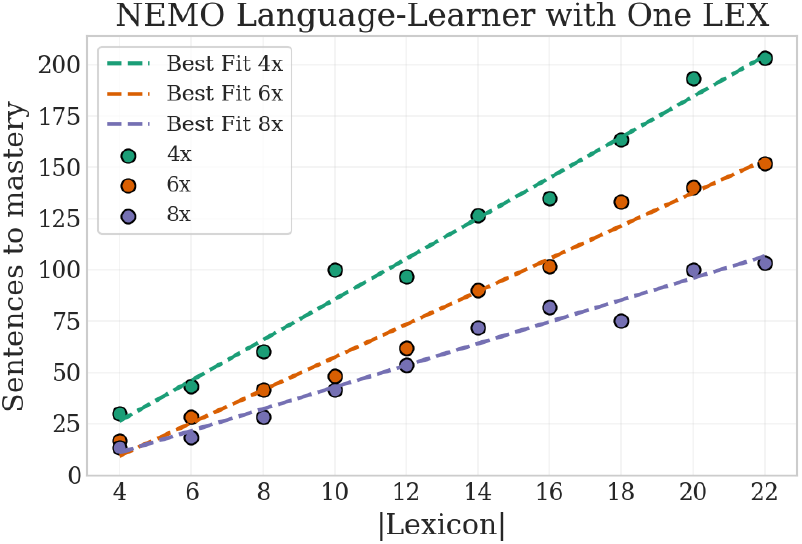
As mentioned in Section 2.3, we implemented a variant of the word learning experiment with *one* LEX area, containing representations of both nouns and verbs. Success is defined as the formation of stable assemblies for each word in the lexicon, such that different assemblies do not overlap too much (less than 50%), and a pathway is formed from assemblies in PHON via LEX to contextual assemblies in VISUAL for nouns and MOTOR for verbs, and also to other semantic areas. The 1-LEX version works only if the value of the cap *k* in PHON is *larger* than those in contextual areas; the same effect can be achieved by increasing the parameter *p* of synaptic density in the fibers between PHON and LEX. The chart plots the number of uniformly-sampled sentences before all words have been successfully learned, for different sizes of PHON assemblies: 4 times those of contextual assemblies (which have *k* = 50), 6 times, and 8 times.

**Figure 7.**
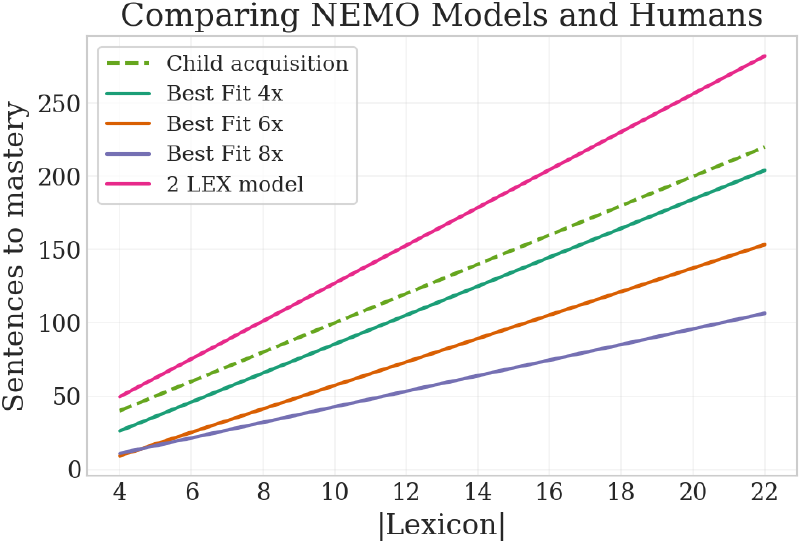
The learning rates (number of uniform sentences to mastery of all words) for the 2 LEX model, and those of the 1 LEX model with varying sizes of PHON assemblies (4 times, 6 times, or 8 times bigger than contextual assemblies) are compared to the predicted child learning rate, based on the state-of-the-art models of (24); for a fully uniform distribution their model predicts 10 ELIs per word, i.e. a line of slope 10 (dashed line). By “tuning” the size of LEX assemblies, or other meta-paremeters of the underlying Nemo model, this idealized child learning rate can be matched. This is one the ways in which the Nemo model can yield predictions of the corresponding parameters in the human brain (see Section 2.3 for further discussion comparing our model to studies of child language acquisition).

Another lesson of language acquisition is that there is a gap between the time a child understands the meaning of a word (determined by cognitive experiments) and the time the child can generate it, see (24). This is also true of our system (see the experiment of Figure 8 in Supplementary Materials). However, in the version with one Lex area understanding and generating seem to arrive together.

**Figure 8.**
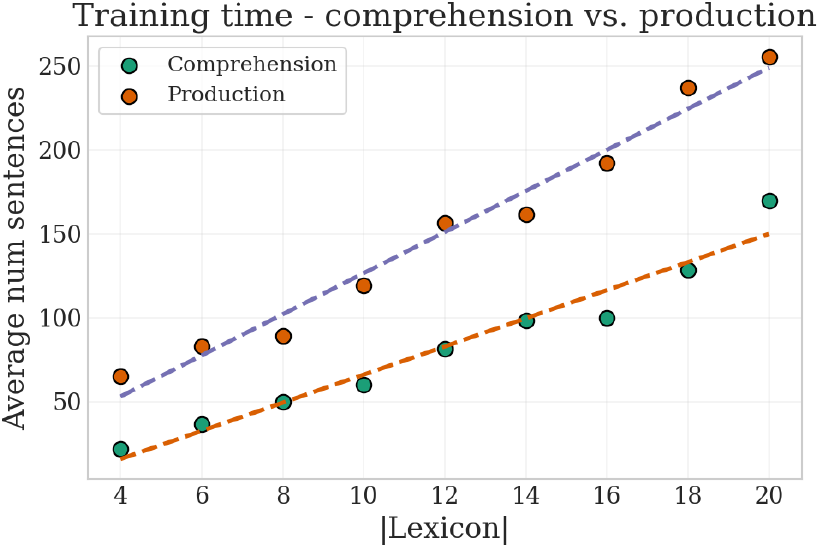
As discussed in Section 2.3, the model learns to *comprehend* words faster than it fully learns them (for generation). We define comprehension as follows: for each word *w*, setting PHON[*w*] to fire, in two time steps (the time it takes to fire into LEX areas and from these into context areas) results in activating the corresponding assembly in either VISUAL or MOTOR (depending on whether *w* is a noun or verb, respectively). Note that this is also a criterion for the full definition of learning (a part of Property 2). Learning times, commensurate with the number of exposures to each word or ELIs, are roughly half of those needed for full learning and generation, in line with findings in child language acquisition (17, 24, 12).

**Figure 9.**
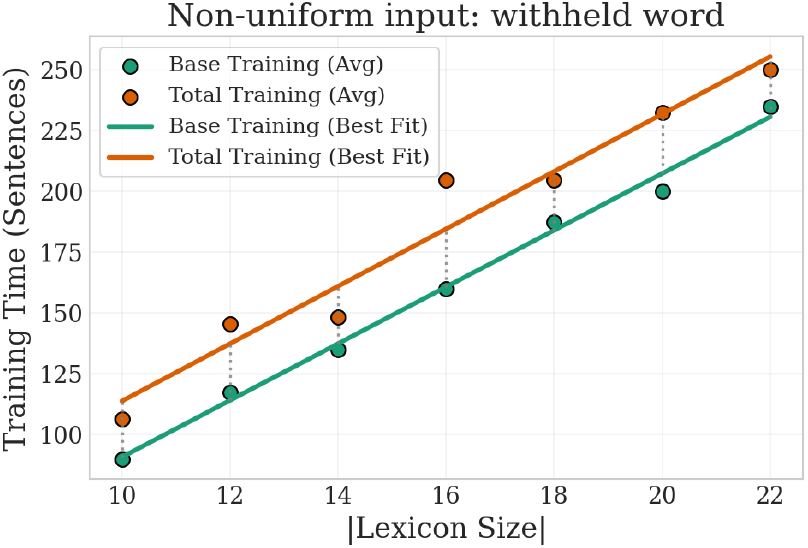
Our system succeeds under various distributions of word and sentence presentations (with uniform sentences being a particularly simple distribution to use to test learning rates). To demonstrate this, this figure and Figure 10 repeat the learning experiment of Section 2 with non-uniform distributions. In this experiment, *n* words (the quantity on the *x*-axis) are presented through sentences. Then, an additional, never-before-seen word is presented, as part of whole sentences with uniform verbs (in the case of a noun) or nouns (in the case of a verb) until that word is fully learned (in the senses of Properties 1-3 as before; note that it is required that all properties continue to hold for all previous words as well). The green line shows training times for the initial *n* words (the same as in the main text Figure 3, and the orange line the *total* training time with the added word. The gap between the lines can be seen as, roughly, the marginal training time for a new word (given memory of *n* previous words), while the uniformly, well-spread out words required about 10 ELIs per word, the additional word added on non-uniformly at the end requires roughly 20 presentations.

**Figure 10.**
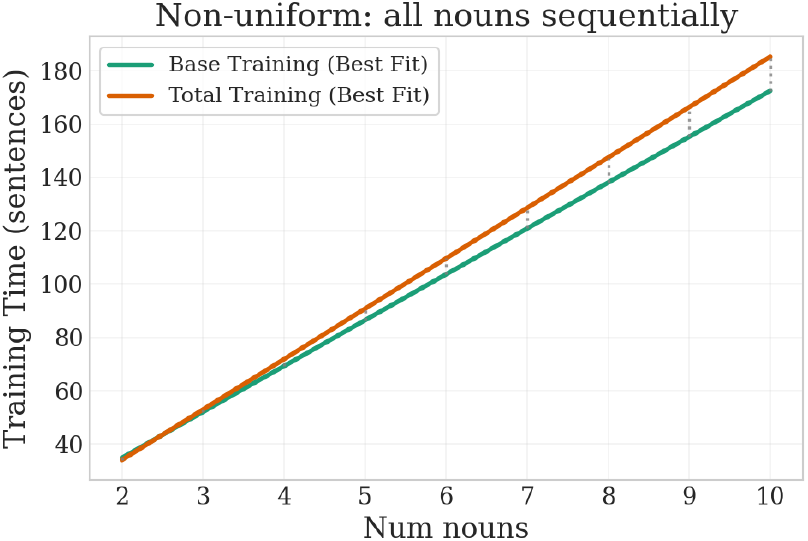
Our system succeeds under various distributions of word and sentence presentations. To demonstrate this, in this figure and in Figure 9 we show the results of repeating the learning experiment of Section 2 with non-uniform distributions. In this experiment, a different number of nouns (*x*-axis) are taught to the system. In an arbitrary order, each noun is presented with uniformly sampled verbs until it is fully mastered (in the sense of Properties 1-3); this is the training time shown in “Base Training” (green line; such forced sequential presentation requires just shy of 20 ELIs per word), substantially more than the spread-out uniform presentations. Sometimes, after the last word is learned, earlier words may fail one of properties 1-3; in this case, earlier words are reviewed (presented again with random verbs, in the same order as originally learned but with one sentence per word, until all words pass all Properties); this total training time is shown in “Total Training” (orange line); the system recovers forgotten words (that have not been seen in a long time) with rather minimal review.

**Figure 11.**
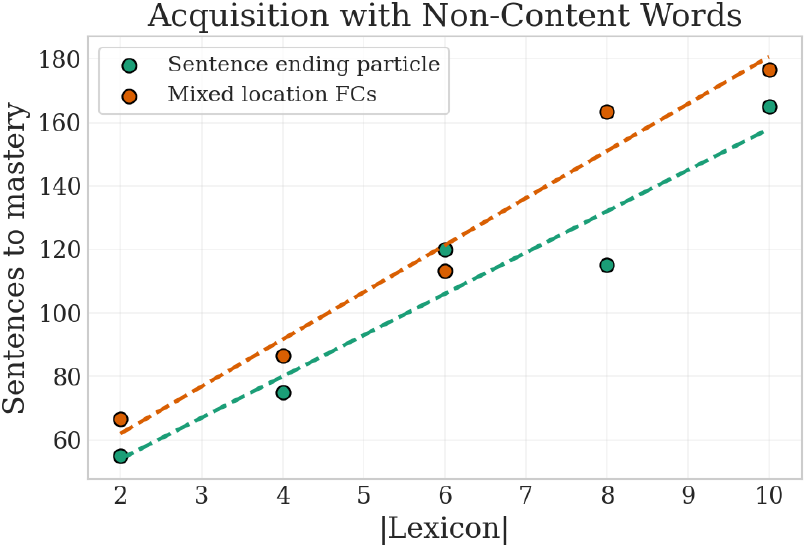
To demonstrate how the NEMO language acquisition system can begin to be extended to broader and more natural settings, we show here that the system succeeds even in the presence of additional, “non-content” words. As in the main system and experiments of Section 2, for increasing sizes of the lexicon, uniform sentences are presented until full mastery of all words in the sense of Properties 1-3. However, PHON now has 5 additional assemblies, corresponding to non-content words. In the “sentence ending particle” experiment, a random non-content word is appended to the end of each training sentence with probability 3*/*4 (that is, the corresponding PHON assembly fires for *τ* time-steps, with all context assemblies firing as usual); training times are shown in green. In the “mixed location FCs” experiment, a random non-content word is inserted at the beginning of the sentence, between noun and verb, at the end of the sentence, or in any 2 of these positions (each template with probability 20%; note in this experiment, a non-content word is included in every training sentence). The addition of non-content words only moderately increases acquisition times for concrete nouns and verbs.

*Child-directed speech (CDS)* is commonly used during language acquisition. CDS favors short sentences, with simpler, usually flat grammatical structure (4) and this finding is cross-linguistic, at least across cultures where CDS is prevalent. In the light of CDS, our input protocol of sentences with 2 or 3 grounded constituents (nouns and verbs), with non-context words sometimes occurring in between, does not seem very unrealistic.

Our system does not cover all aspects of word acquisition. It is best seen as a model of the period extending up to about 30 months where word-learning is driven by ELI exposure; indeed, experimental evidence largely matches our models’ predictions. After this period, during what is often called the “vocabulary explosion,” other mechanisms (such as perhaps learning basic syntax and instrumentalizing that and functional categories to immediately classify words by their grammatical properties) may become more important.

### 2.4 Syntax: learning the word order, and learning to generate sentences

Next we turn to *syntax*, and, of all the interrelated linguistic phenomena entailed by syntax, we focus on the most central: learning the language’s basic *word order*. We test that the system can *produce novel sentences* when appropriate sensorimotor stimuli are presented — for example, articulate the sentence “Sister pushes Dad” upon sensory input of the corresponding scene, even if this scene was never seen or described before.

#### Word orders and grammatical moods

Languages can be broadly classified as belonging to one of six basic categories based on the default ordering of the sentence’s subject, object, and verb, known as the *constituent order* or *word order*. For instance, English is an SVO (subject-verb-object) language and so is French, while Japanese is SOV and Tagalog is VSO.

Let us note here two apparent linguistic universals (7): (a) Word orders starting with O are extremely rare; and (b) in attested languages with a rigid or default word order, the intransitive order is almost always the same ordering with O omitted^1^ — for example, if the default transitive order is SOV, the intransitive order is SV. As we shall see, these two linguistic facts *can be predicted by the workings of our device*.

We also note that many languages have varying constituent orders depending on semantic factors such as *mood* — whether the sentence is a statement, question, command, etc. —, syntactic factors (e.g., as part of a dependent clause), or just as a result of emphasis and pragmatics (especially in languages with relatively free word order such as Polish). While we do not set out to handle the entire scope of these complexities, we will show that our model can acquire and output different word orders for different *grammatical moods* (declarative, interrogative, subjunctive, etc.) via a simple mechanism; we believe that this mechanism can be extended to model syntactic and pragmatic variations of the word order.

#### Thematic roles

Analyzing a complex scene by assigning thematic roles to the participants of the scene — determining who is doing what to whom, and who is watching — is an important skill for animal survival. Cognitive experiments, e.g. (14**?** ) suggest that there is a computation of roles in a scene based on visual cues.The precise seat of the non-linguistic mechanism that assigns roles to perceived agents in a scene remains unknown, but a related kind of computation takes place in the left temporoparietal junction (lTPJ), believed to be a multimodal hub integrating information from diverse perceptual pathways to “create coherent, spatiotemporal representations of the complex dynamic situations” (3). We hypothesize that there is a system in the primate brain, perhaps in communication with the TPJ, which *computes thematic roles of scenes*. Our Nemo implementation (Figure 4) contains three role areas, Role_agent_, Role_action_, and Role_patient_, as well as a fourth area, Role_scene_ synaptically connected to all three. We assume that, when a scene is presented, such as a dog chasing a cat, there are assemblies in the three role areas Role_agent_, Role_action_, and Role_patient_ synaptically connected to the assemblies for “dog,” “chase,” and “cat” in Lex_1_ and Lex_2_ while Role_scene_ contains an assembly for this scene synaptically connected to the three.

**The full language system** has several areas beyond those in Figure 2: (a) the four Role *areas:* Role_agent_, Role_patient_, Role_action_, and Role_scene_; (b) three *syntactic* areas, Subj, Obj and Verb, and (c) a Mood area, plus 18 fibers conneccting them as shown in Figure 4. Finally, the three Role areas, and separately the three syntactic areas, are in *mutual inhibition* — the only use of this feature of Nemo in this model.

### 2.5 The syntax learning experiment

Following the word learning experiment of Section 2.1, we input grounded sentences, both transitive and intransitive to the full system. Grounding now means that, with each input sentence, e.g. “Sister grabs the toy,” a scene to the same effect has been input to the sensory system, and its thematic roles (patient, agent, action) are represented as assemblies in the Role areas.

For each consecutive word in each sentence, the corresponding assembly in Phon is fired for *τ* time steps, allowing firing to propagate throughout the lexical areas, Role areas, and finally into the syntactic areas. The constituent order is learned through plasticity in the fibers between the syntactic areas and the Role areas.

#### The experiment

We fix a language, a number of moods, and for each mood, one of the six possible constituent orders. We input a sequence of random transitive and intransitive sentences for randomly chosen moods in the correct order. Every 10 sentences, we *probe* by running a generation experiment: we test, upon the presentation of a randomly generated novel scene (e.g., a dog eating a cookie, presented as three assemblies in the three Role areas), whether the device will generate the appropriate sentence with the correct word order: “the dog eats the cookie” (the details of generation are presented next). Success of the experiment for this mood and word order is defined as the successful generation of a transitive sentence and an intransitive sentence, both sampled randomly and withheld during training.

#### Generation

The human brain has the ability and means to *not* generate the sentence “the dog runs” even though the dog may be running. Hence generation must involve a prefrontal cortex assembly, called *trigger*, which implements the decision to generate. A scene (imagined or seen) is presented in the Role areas, and the intended mood is determined by an assembly in the Mood area. The generation algorithm entails simply firing the trigger assembly.

### 2.6 Results of the experiment on word order and generation

The results of our experiments are shown in Figure 5. The device succeeds in learning the constituent order, for several languages with different word orders and for a varying number of moods, and the number of sentence presentations that is needed for this appears to grow linearly with the number of moods. The six word orders differ substantially in difficulty, and our results (Subfigure 5c) are in striking agreement with what linguists have known for decades (7): The word orders OSV and OVS are exceedingly rare, and VOS is a close third. These are precisely the three word orders that are the hardest to learn by our system: If the human language system were structured like the one presented here, the relative frequency of word orders would be predicted by our model. Also, in about 99% of languages the word order in intransitive sentences is the same as the word order in transitive sentences with O omitted — and this fact is again reflected in our model’s difficulty of learning word orders.

## 3 Discussion

We presented what we believe is the first biologically plausible system that can learn the rudiments of natural language — lexicon as well as basic syntax and semantics — through the presentation of a modest amount of grounded sentences in any language. In a first phase, our system learns the meaning of tangible words, and subsequently in a second phase syntax is learned qua the word order of the language. Although the resulting system does not begin to exhaust the range of human language, we believe that it is of some interest. Were our system’s linguistic abilities to be detected in a nonhuman animal, the prevailing views about the exclusivity of language would have to be revised.

Many facets of language are not covered by our system: The learning of speech sounds and phonology has been bypassed, while adjectives and functional categories are not considered. We believe that many of these missing aspects can be accommodated by more or less simple extensions of the present system. *Abstract words* — words that are not directly grounded in sensorimotor experience, such as “bravery,” “available,” and “negotiate” — are an important next step that will require, among others, the development of a representation scheme for such words and the introduction of new semantic brain areas — the representation of abstract words is a well known open problem in cognitive linguistics, see for example (2). It would also be very intriguing to experiment with interactions between Nemo artifacts and large language models (LLMs) as a means to help the former grow in more demanding directions such as the quintessential *social* aspects of language. Beyond language, *reasoning* in its various genres — logical, heuristic, categorical, by analogy, etc. – is an important next challenge for Nemo artifacts.

Our system can, however, be extended to handle *multilinguality:* A new area can be added containing an assembly for each language known to the speaker, very much like the representations of grammatical moods in the Mood area. This assembly would be active while learning, or using, the corresponding language. Initial experimentation with this idea is encouraging, but more work is needed.

We believe that our language acquisition system is biologically plausible not only because it is strictly based on the Nemo framework, but also because its *tabula rasa* brain architecture — the areas and the fibers connecting them — can be plausibly delivered by mammalian developmental processes. One possible exception are the assemblies in the Mood area, each representing a different mood of the language. It can be argued that such a mood assembly can be created during learning on the basis of various sensory and mental-state cues associated with the mood (e.g., the tension in the discourse and the special tone of voice when a question is asked). However, more work is needed to fully justify this point.

In addition, our treatment ignores the *hierarchical* structure of language, believed by many to be of paramount importance: a sentence may have another sentence embedded in it, and that sentence as well, and so on. For example, the utterance *“Melville, who I love, wrote a book about whaling that became a classic”* contains three sentences. Our language system can be extended very naturally to record the *syntax tree* of complex utterances, also through synaptic strengths, by introducing new areas corresponding to Broca’s, see (10), and (22) for a Nemo-based system for handling embedded sentences. In fact, hierarchical structure may actually be *needed* for our system to treat language in all its complexity, as it has been observed in previous work (8) that there *is* a limit to the length of chains of assemblies in an area that a Nemo system can maintain reliably.

Notwithstanding the important and challenging research directions that lie ahead and the limitations mentioned above, the artificial system for basic language acquisition based on Nemo we present here is the first of its kind and, in addition to its value as a proof of concept, it may be of some use to neurolinguistic research. Its design decisions are intended to largely reflect current thinking, and hence the resulting artifact is a concrete hypothesis that can possibly help guide further research on the human language system through the generation of testable predictions and the deployment of more advanced versions. We conclude with several such predictions – what one should expect if the human language system were to be structured as the Nemo artifact described in this paper:

- The syntactic phase of our system requires a dense network of dozens of fibers connecting the seven brain areas associated with syntax (recall Figure 4), an extraordinarily complex neural machinery which seems evolutionarily and developmentally demanding. It is known that the arcuate fasciculus bundle of fibers is more extensive in the language-dominant hemisphere in humans, an asymmetry that is unique among mammals and an evolutionarily novel trait (11). Certain tracks of this bundle have been found to be myelinated after the first year of life (10). We hypothesize that these facts are not unrelated to the advent of syntax in the fourth semester of human life.
- We found that those word orders that are harder to learn by our syntax acquisition system are those that are most rare among world languages. We hypothesize that this is not coincidence, but that the same word orders are the hardest to learn also for the human language system, and as a result they present an evolutionary burden that renders such languages rare; languages are known to evolve and the word order is one of the characteristics that can change, as witnessed by modern Greek, Hebrew, and Italian, whose ancestral languages had different word orders. Our syntax experiment also predicts that learning of the three rarest orders OSV, OVS and VOS requires the presentation of substantially more transitive sentences than intransitive.
- Concerning the question of whether verbs and nouns are represented in the same or separate brain areas, our results are equivocal. In the model with the same brain area for both nouns and verbs, learning is markedly more efficient than the alternative, and this efficiency seems to increase with the amount of synaptic connectivity between PHON and this area; however, the single LEX area model (a) requires several times larger assembly size and/or more connectivity than the two-area model, and (b) it fails to reproduce the fact that, in human language acquisition, the ability to understand the meaning of a word precedes somewhat the ability to generate the word, whereas the model with two areas confirms this fact.

## 4 Materials and Methods

### NEMO Simulations

All simulations in this paper were built upon the open source NEMO Python Library developed by Mitropolsky and Google (23). This library is an exact implementation of the NEMO dynamical system as presented in the legend Figure 1 and in Subsection 4 below. The firing operation is implemented using a *lazy-evaluation* trick that speeds up computation (this heuristic is described in older NEMO work such as the Appendix of (27)).

For the word learning experiments of Section 2.1, the data presented in (a) and (c) of Figure 2 was generated using NEMO parameters: *p* = 0.05, *n* = 100, 000, *β* = 0.06, and *k* = 50 in Lex_1_, Lex_2_ and 100 elsewhere. For each lexicon size (or number of verbs in (c)), the experiment is repeated for 5 independent repetitions and averaged. To demonstrate robustness relative to a reasonable range of parameters, we repeat the experiment of (a), only for lexicon sizes 10 and 20, with a variety of other parameter settings and verified success in finite time: all combinations of *n* = 10^5^, 10^6^, *p* = 0.05, 0.02, 0.01, *β* = 0.1, 0.06, 0.05, 0.03 were verified.

For the constituent order experiments of Section 2.5 and the data in Figure 5 (b) and (c), we used NEMO parameters *p* = 0.05, *n* = 100, 000, *β* = 0.06, and *k* = 50 in Lex_1_, Lex_2_ and 100 elsewhere. Each experiment (number of moods, or constituent order setting) was averaged over 10 independent repetitions. The default ratio of transitive: intransitive sentences used was 70%, but 30% in experiments labeled as *<* 50% transitive.

## Code Availability

Our custom Python library is publicly available online for open use under the MIT license (21). The library includes the exact commands used for the learning experiments of Sections 2 and 3, and the output data is summarized in Figures 3 and 5.

### Details of the NEMO model

The caption of Figure 1 presents a condensed summary of the Nemo model. The full details, in particular of the version used in this paper which innovates on the original model of (27) in several ways, are as follows:

An instance of Nemo so far is specified by:

- A set of brain areas;
- The number *n* of excitatory neurons per brain area
- A set of ordered pairs of brain areas that are connected through fibers;
- The connectivity probabilities *p*
- the size of caps in each area; while the caption of Figure **??** simplifies the presentation slightly, we allow a different cap *k*_*A*_ for each area *A*).
- The plasticity coefficient *β*
- The list of sets of areas in mutual inhibition.

Each instance of Nemo defines a dynamical system, where the *state* at time *t* consists of (a) for each neuron *i* a bit 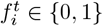 denoting whether or not it *fires* at time *t*, and (b) the synaptic weights 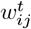 of all synapses in the model. Given this state at time *t*, the state at time *t* + 1 is computed as follows:

1. For each neuron *i* compute its *synaptic input* 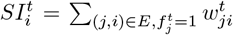, that is, the sum total of all weights from pre-synaptic neurons that fired at time *t*.
2. Next we compute 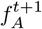, a boolean indicating whether there is any firing in *A* at time *t* + 1. By default,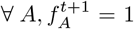. For each set *S* = *A*_1_, …, *A*_|*S*|_ of areas under mutual inhibition, compute the total synaptic input for each area 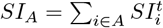. For *A*_*i*_ ∈ *S*, if 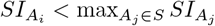 set 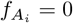 (breaking any ties arbitrarily).
3. For each neuron *i* in area 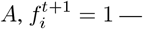 that is, *i* fires at time *t* + 1 — if *f*_*A*_ = 1 *and i* is among the *k* neurons *in its area* with the highest 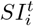 (breaking any ties arbitrarily).
4. For each synapse (*i, j*) *∈ E*, 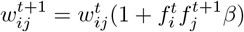 ; that is, a synaptic weight increases by a factor of 1 + *β* if and only if the post-synaptic neuron fires at time *t* + 1 and the pre-synaptic neuron had fired at time *t*.

## Acknowledgments

The authors thank Angela Friederici and Evelina Fedorenko for helpful discussion and feedback on the results in this paper.

## Data and materials availability

Our custom Python library is publicly available online for open use under the MIT license (16). The library includes the exact commands used for the learning experiments of Sections 2 and 3, and the output data is summarized in Figures 3 and 5. All data that is output by these experiments is available in the main text figures.

## 5 Supplementary Materials

### 5.1 Additional Experiments and Extensions

## Footnotes

1 Based off the World Atlas of Language Structures Online (7), out of 1185 languages with default transitive and intransitive orders recorded, we found at most 11 that *may* invert *S* and *V* in intransitive order relative to transitive order.

